# Dynamics of infection and immunity over 50 years as marine stickleback adapt to freshwater

**DOI:** 10.1101/2025.02.04.636402

**Authors:** Pranav Sriramulu, Dolph Schluter, Daniel I. Bolnick

## Abstract

When a species colonizes a new environment, it may encounter new parasites to which its immune system is poorly adapted. After an initial spike in infection rates in the naïve founder population, the host may subsequently evolve increased immunity thereby reducing infection rates. Here, we present an example of this eco-evolutionary process, in a population of threespine stickleback (*Gasterosteus aculeatus*) that was founded in Heisholt Quarry, a man-made quarry pond, in 1967. Marine stickleback rarely encounter *Schistocephalus solidus* tapeworms (which require freshwater to hatch), and so remain highly susceptible to infection. Initially, introduced marine fish were heavily infected by *S.solidus*. They exhibited low levels of fibrosis, a heritable immune trait which some genotypes activate in response to infection, thereby suppressing tapeworm growth and viability. By the 1990’s, the Heisholt Quarry population exhibited high rates of fibrosis, which partly suppressed *S.solidus* infection. This increased immune response led to reduced infection rates and the tapeworm was apparently extirpated by 2021. Because fibrosis has a strong genetic basis in other stickleback populations, we infer that the newly founded stickleback-parasite interaction exhibit an eco-evolutionary process of increased immunity that effectively reduced infection. The infection and immune dynamics documented here closely match those expected from a simple eco-evo dynamic model presented here.

**IMPACT SUMMARY:** Parasite-host relationships are a great framework within which to study the mechanisms driving evolutionary change, both in terms of ecological dynamics, selection pressures, and the phenotypic basis of adaptation to environmental change. One classic example of adaptation is the loss of armor plating as marine stickleback fish colonize freshwater habitats. However, the parasitological and immunological adaptations in new freshwater populations are less clear. When marine fish colonize freshwater they encounter an unfamiliar tapeworm parasite, *Schistocephalus solidus*, to which they are poorly adapted. Many (but not all) freshwater populations evolve an inducible fibrosis response to *S.solidus*, a build-up of scar tissue that limits parasite growth and survival. However, we did not know the time frame over which marine fish, invading freshwater, will evolve this immune trait. To resolve this, we tracked the eco-evolutionary dynamics of *Schistocephalus solidus* tapeworm infection and the fibrosis immune response over 50 years, in an artificial quarry pond populated by susceptible marine fish in 1967. We show that infection rates were initially high, as the marine fish encountered a freshwater parasite to which they were not adapted. Fibrosis rates subsequently increased, reducing the infection rates and by 2021 *S.solidus* was apparently eliminated. This process matches a simple eco-evolutionary model of an inducible immune response, and provides a clear example of how hosts can adapt to and suppress novel parasites. Although fibrosis is typically viewed as an immune pathology contributing to disease in humans, we show that in stickleback it has a key adaptive value in protection against helminth infection.

## INTRODUCTION

The infection rate of a parasite in a host population depends on both the parasites’ ability to enter and establish within hosts, and hosts’ ability to resist infection. Both species often exhibit genetic variation in their respective traits controlling infection (Thompson and Lymbery 1996 ), and these traits can be subject to strong selection (Crossan et al. 2007, Ebert 2008, Lachish et al. 2011). Therefore, the infection rate can change as one (or both) species evolve to adapt to each other. This adaptation may be particularly important in newly founded new host (or parasite) populations (Nørgaard et al. 2019). When hosts disperse into previously unoccupied habitats, they may leave behind parasites from their native range (the ‘enemy release hypothesis’), but may also encounter novel parasite species (or genotypes). Hosts are likely to be susceptible to any newly acquired parasite, lacking a history of adaptation to it. The resulting high infection rates may impose strong selection for greater host immunity. We therefore expect to observe rapid evolution of immune resistance in newly founded host populations. For example, Bonneaud et al. (2011) found that house finches from an area with a longer history of exposure to an introduced bacterial pathogen causing conjunctivitis were less affected than finches from recently exposed populations. Such host adaptation should lead to declining parasite prevalences over time. However, there are few examples where researchers have simultaneously tracked host immune evolution, and the resulting changes in parasite prevalence. To track such eco-evolutionary dynamics, one ideally needs to monitor both infection rates and host immune traits through time, in a newly founded population. Here, we present an example of this eco-evolutionary dynamics, taking advantage of museum collections from an experimentally founded population of threespine stickleback established in the late 1960’s.

Threespine stickleback (*Gasterosteus aculeatus*) are a predominantly marine species of fish, but they frequently established permanent freshwater populations in north temperate coastal lakes and streams following Pleistocene glacial retreat. These freshwater populations have evolved a variety of adaptations to their new environmental conditions (eg. salinity, predators, diet, parasites). Most famously, marine stickleback have extensive lateral armor plates, but freshwater populations evolve reduced armor coverage (Colosimo et al. 2005). This armor loss is repeated in many independent freshwater stickleback populations around the northern hemisphere. This adaptation can be quite rapid: in one example an anthropogenically founded population in Loberg, Alaska evolved from having nearly the entire population being high-armored to only 11% of the population being high-armored, in only 12 years (Bell et al. 2004).

When stickleback colonize freshwater, they come into contact with a variety of parasite species not typically found in their ancestral marine environment. An especially common example is the cestode *Schistocephalus solidus*, a tapeworm specialist on threespine stickleback that can grow to nearly half the fish’s body mass (Nordeide & Matos 2016). *S. solidus* eggs hatch in freshwater, producing coracidia larvae which are ingested by several genera of cyclopoid copepods. If an infected copepod is ingested by stickleback, the tapeworm penetrates the intestinal wall and matures in the fish’s body cavity. *S.solidus* reproduces in piscivorous birds such as loons (Barber and Scharsack 2009), which disperse the parasite between freshwater habitats. The tapeworm’s eggs hatch poorly in brackish water, so *S. solidus* is rare or absent in marine and anadromous stickleback. Without regular exposure to *S.solidus*, the marine populations have not evolved resistance and are readily infected in laboratory experiments (Weber et al. 2017a). Thus, newly established freshwater stickleback encounter *S. solidus* tapeworms that their marine ancestors have not evolved to resist (Weber et al. 2017a). For instance, Cheney Lake (Alaska) was repopulated by stickleback in the 1990s after the lake was poisoned to remove northern pike (Bell et al. 2004), and the population currently exhibits nearly 100% infection rates (Bolnick et al. 2024a). Similarly, when anadromous fish do migrate into lakes with resident freshwater populations, they are often heavily infected (Confer et al. 2012).

In long-established lake populations (likely colonized ∼12,000 years ago), stickleback have evolved increased resistance to *S. solidus*, relative to marine relatives (Weber et al. 2017a). In some lakes, this resistance includes a peritoneal fibrosis response to *S. solidus* antigens (Weber et al. 2022; Hund et al. 2021). Fibrosis involves the creation of thick layers of collagen, which will then act as an adhesive force between various bodily organs (Weber et al. 2022). This fibrosis is similar to an immune pathology associated with various inflammatory responses in humans (Turola et al. 2015), and can be induced by immune stimulants across the diversity of jawed fishes (Vrtilek and Bolnick 2021). Despite this ancient origin of fibrosis, not all freshwater stickleback populations initiate fibrosis in response to *S.solidus*. The ability to initiate fibrosis in response to *S. solidus* antigens has been shown to be heritable and mapped to a single locus of large effect containing the fibroblast regulatory gene Spi1b (Weber et al. 2022). Importantly, Spi1b is on linkage group II, unlinked from the quantitative trait loci typically associated with adaptation to freshwater (e.g., linkage group IV).

The evolution of fibrosis has been inferred by comparative studies of phenotypic and genetic differences among long-established freshwater populations, and marine populations (Weber et al. 2022, Hund et al. 2021). Modern marine stickleback (a proxy for the genotypes of ancestral marine stickleback; Kirch et al. 2021) do not initiate a fibrosis response to cestodes, or cestode protein injections (Hund et al. 2021). This lack of fibrosis response is consistent with the rarity of tapeworm infections in marine fish, suggesting a lack of evolutionary pressure for fibrosis-based immunity. In contrast, many freshwater populations (established ∼12,000 years ago) do exhibit a fibrosis response to infection in numerous lakes in both British Columbia and Alaska (Weber et al. 2022). The presence of fibrosis in many independently evolved freshwater populations suggests strong selection to gain fibrosis-associated immunity, following colonization of freshwater ecosystems. Although fibrosis response is apparently gained during freshwater colonization, some freshwater lake populations subsequently exhibit positive selection on Spi1b deletions that suppress fibrosis (Weber et al. 2022). This reversion to a non-fibrotic strategy reflects fitness costs imposed by the fibrosis pathology, reducing feeding, breeding, and mobility (De Lisle and Bolnick 2021, Matthews et al. 2023).

Inferences about fibrosis evolution would be much stronger if we could directly document changes in fibrosis during the early stages of freshwater colonization. This is possible, because a number of researchers have experimentally founded freshwater populations of stickleback by transplanting marine fish into fishless freshwater habitats (e.g., Bell et al. 2016; Aguirre et al. 2021). For instance, in 1967, Don McPhail introduced marine stickleback into a fishless pond in the former Heisholt Quarry. Note that *S. solidus* may be readily distributed into such fishless lakes by passing piscivorous birds such as loons, mergansers, kingfishers, and herons, the terminal host for *S. solidus*. Therefore, the parasite may be present in the copepod community even before the obligatory secondary host is introduced.

Based on recent findings described above, we make several testable predictions about the eco-evolutionary dynamics of fibrosis and infection in such experimentally founded populations. First, we expected that the early generations of stickleback to colonize freshwater lakes will have high cestode infection rates. When the immunologically naïve marine genotypes first encounter freshwater *S. solidus*, they will mostly lack genotypes needed for effective resistance. Despite high infection rates, we expected to see little infection-induced fibrosis in these early generations, as marine fish mostly lack pro-fibrosis genotypes (Vrtilek and Bolnick 2021, Hund et al. 2022). Second, if any colonists carried genotypes with the capacity to initiate fibrosis in response to *S. solidus* exposure, they should be better able to suppress infection and gain a fitness benefit. Therefore we expect increased fibrosis across generations as genotypes capable of responding to infection become more common. Third, we expect that this increasing fibrosis response would suppress cestode prevalence. Finally, the decline in *S. solidus* abundance may then lead to a reduction in visible fibrosis, because the fish are no longer being exposed and triggered to initiate fibrosis. We begin by illustrating these hypotheses with a simple eco-evolutionary model (Fig. 1). To test these hypotheses, we measured changes in infection rates and fibrosis using a 50-year series of museum samples from Heisholt Quarry.

**Figure 1.**
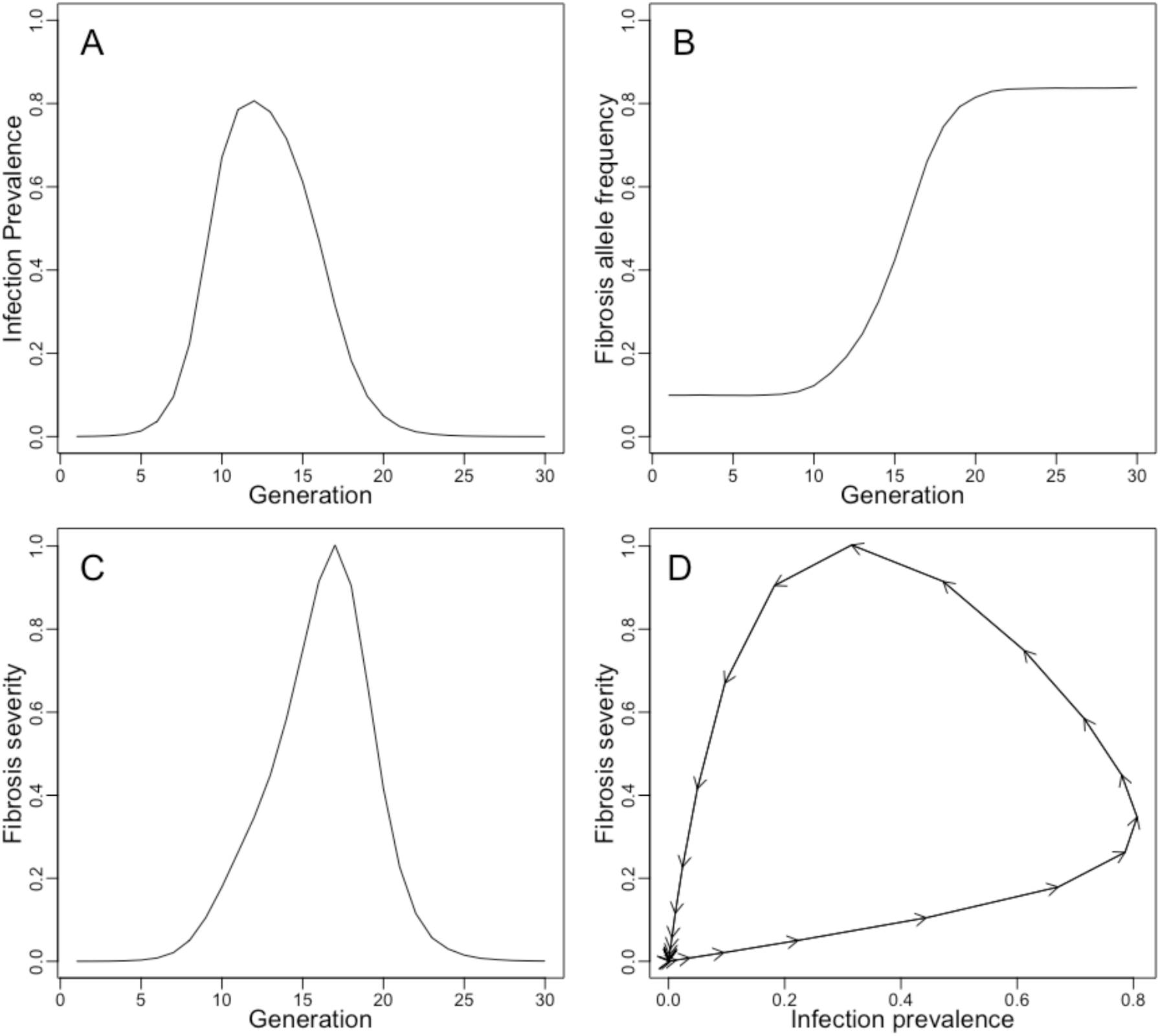
A simple model of eco-evo-immunodynamics in a newly colonized host population exposed to a novel parasite, which triggers an inducible immune response in one host genotype. Panel (A) shows dynamics of parasite prevalence (the proportion of fish with 1 or more parasites) starting with L = 100 infectious larvae in a lake with V = 500,000, and reproductive rate λ =15 by surviving parasites. Panel (B) shows host evolution, tracking changes in the frequency of resistance allele R, starting with a founding population frequency *P(R)* = 0.1, assuming fitness costs of the immune response *c = 0.15,* and costs of infection *i = 0.8.* The inducible immune phenotype (C) shows an initial increase as allele R becomes common, and a decline as parasite exposures become scarce. The resulting eco-evolutionary dynamics (D) lead to a cyclic arc through infection-trait space over 50 generations, though the host population genotypes are greatly changed from the start to the end. Code for the simulations is available in the data repository for this paper, on Figshare DOI: 10.6084/m9.figshare.28342601.

### An eco-evolutionary model of an inducible immune defense

We used a stochastic individual based simulation to model the population dynamics of a parasite in a newly founded population of initially susceptible hosts. We also tracked the population genetic dynamics of two alleles at a single locus: allele *R* confers resistance to infection through an inducible immune response (e.g., fibrosis), whereas allele *r* fails to respond to infection and is susceptible. Specifically, genotype *rr* has a fibrosis value of 0 regardless of infection status. Genotype *Rr* has a fibrosis value of 0 if unexposed to the parasite, but responds to infection by increasing fibrosis to 2. Genotype *RR* also has a baseline value of 0 but responds to infection with a fibrosis value of 4. We did not explicitly model host population dynamics, assuming hosts are at a stable carrying capacity set by other ecological factors.

We simulated a stable population of 100,000 host individuals, founded with a small initial allele frequency *P(R)*. Genotypes are randomly assigned to individuals from a binomial random number generator with two draws per host with expectation *P(R)*. There was an initial input of *L* infectious parasite larvae (e.g., procercoids in copepods) diluted into a lake volume *V*. The density of larvae in the water dictates fishes’ exposure rate (*E = L/V*). Each fish is exposed to a random number of tapeworms, drawn from a Poisson distribution with expectation *E*. After exposure, fish initiate fibrosis to a level set by their genotype. Parasites survival is determined by a binomial random draw, with probabilities 1, 0.5, and 0 in fish with fibrosis scores of 0, 2, and 4 respectively. Fish retain fibrosis even if their parasite is eliminated. Surviving parasites reproduce, each contributing λ new infectious-stage larvae that are diluted into the lake. We omit details of modeling transmission through other host species (e.g., copepods and birds for *S.solidus*). Host fitness is set to *w* = 1 for an uninfected individual, *w* = 1-*c* for an individual with fibrosis that eliminated the infection (to account for costs of fibrosis), *w* = 1-*i* for an infected fish without fibrosis, and *w* = 1-*(c+i*) for an individual with both fibrosis and a surviving infection. Each genotype’s mean fitness (*w̄*) was calculated by averaging across all individuals (with/without fibrosis, with/without infections). The next generation’s allele frequency is set by the standard population genetic formula based on each genotype’s frequency *F* and mean fitness:

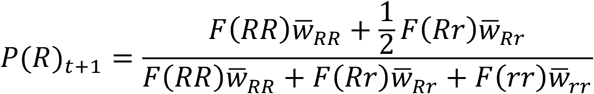

We then generated a new generation of host genotypes drawing from this allele frequency, and repeated the steps of exposure, fibrosis, and reproduction. We present an example of the dynamics resulting from this model, to illustrate a possible outcome, but do not here present an exhaustive search of parameter space.

Initially the parasite population size is small, but quickly grows (Fig. 1A) thanks to an abundance of susceptible hosts with genotype rr. As the parasite population grows, selection favors resistant allele *R,* which becomes more common (Fig. 1B). Because of this evolution, we observe a transient increase in the inducible immune trait (Fig. 1C), as resistant genotypes are common and infections still abundant. However, increased immunity reduces parasite fitness, and eventually the parasite is eliminated. As the parasite declines the fibrosis trait declines as well, since it is no longer being induced, even though the allele for fibrosis persists at high frequency. The result is non-linear cyclic covariation between infection prevalence and the fibrosis phenotype (Fig. 1D).

## METHODS

### Field sampling

In 1966, Heisholt Quarry on Texada Island (latitude 49.772 N, longitude -124.596 W) ceased commercial operation and was flooded to create a small pond comprising of two basins connected by a seasonal stream (MacLean 1974). Around two thousand marine stickleback were introduced in 1967 from a nearby coastal stream on the island (MacLean 1974, Maclean 1980).

Stickleback were sampled from Heisholt Quarry in 1974 by Don McPhail, in 1996, 1997, 2000, and 2005 by Steve Vamosi (Vamosi 2006), and in 2021 by D. Schluter, Stephanie Blain, and Mackenzie Kinney. Stickleback were collected by setting minnow traps in Heisholt Quarry. The traps were set along the lake bottom at 1-2 meters depth. After capture, fish were euthanized with MS-222 then preserved in formalin (10% neutral buffered) and then stained in Alizarine Red before storage in 70% isopropanol. With the exception of the initial sample (which predated current regulations), sampling was done with approval of the government of British Columbia and Animal Use and Care approval from the University of British Columbia (UBC Animal Care permit A20-0050; SU21-618979 for 2021).

### Dissection

We measured fish mass (grams) and standard length from the head to the end of the caudal peduncle (mm). We counted the number of stained armor plates on each side of the fish before beginning dissection. Using scissors we cut from the cloaca along the ventral surface, through the pectoral girdle up to the hyoid, and spread the body cavity open to score fibrosis, moving the organs with forceps to detect fibrotic adhesions between organs. Fibrosis was scored on a scale of 0-4 developed by Hund et al. (2022). In fish with no fibrosis (score of 0), the liver, intestine, and gonads move freely and separately from each other and from the body wall. In slightly fibrotic fish (score of 1), there are fibrotic threads between organs that tear easily. In moderately fibrotic fish (2) the liver and intestines are usually strongly connected, requiring force to separate. In highly fibrotic fish (3), the organs are attached to the body cavity by fibrotic threads. In extremely fibrotic fish (4), the adhesions are extensive enough that it is difficult to pry open the body wall and the organs may tear in the process. These ordinal fibrosis scores are highly repeatable between observers (Bolnick et al. 2024). We inspected the body cavity for *S.solidus* cestodes, and counted and weighed them if present. We also determined the fish’s sex during the dissection by inspecting gonads. Sample sizes per year were n = 56 in 1974, n=31 in 1996, n=53 in 1997, n=28 in 1999, n=56 in 2000, n=17 in 2005, and n=27 in 2021.

### Data analysis

Marine stickleback are well known to adapt to freshwater by evolving reduced armor plating. To check whether this was consistent for the introduced Heisholt Quarry population, we used linear regression to look at the change in time of armor plating (as an average of the left and right sides). Because the armor plating trend through time was highly nonlinear, we used the *nls2* package in R to fit an exponential decay function (mean plate number ∼ *a* + *b*(e^(-*c*\*year)^). We also used regressions to test for temporal changes in log length and log mass. All data analysis was done in R (R Core Team 2023). The data files and R code are archived on a public repository (Figshare DOI: 10.6084/m9.figshare.28342601).

To examine the relationship of in infection rate over time, we used a binomial generalized linear model (GLM) to test whether the prevalence of *S.solidus* infections changed directionally over time. Fibrosis frequently results in suppressed tapeworm mass, so we tested whether log cestode mass varied as a function of year, infection intensity, and fish mass (using stepwise Akaike Information Criterion (AIC) to select the model with best fit.

We expected an increase in fibrosis over time as stickleback evolved the capacity to initiate fibrosis in response to tapeworm exposure (Fig. 1). However, we also anticipated that this increase might be transient, if fibrosis reduced tapeworm exposure rates enough so fish were no longer stimulated. We therefore used a quadratic regression to test for non-monotonic temporal changes in mean fibrosis across years. We also expected to find an association between mean fibrosis and infection metrics (intensity, or log mass), given past evidence that fibrosis suppresses infection (Hund et al. 2022).

If the fibrosis response is purely plastic and did not evolve over time, the relationship between individual infection status and fibrosis should be invariant over time. Conversely, if stickleback evolved an increased fibrosis response to infection, we should find that individual fish fibrosis depends on an infection*year interaction (or conversely, a fibrosis*year interaction on infection presence/absence).

## RESULTS

Consistent with previous studies of stickleback freshwater colonization (Bell 2004, Colosimo et al. 2005), we observed changes in body size and armor in the first few decades after marine stickleback were introduced to Heisholt Quarry. By 1974, several years after the initial introduction, stickleback were already polymorphic for armor number and on average partly plated. Fifteen individuals were fully plated (20-35 plates per side), 13 were partially plated (10-20 plates per side) and 17 were low plated (< 10 plates per side). By the late 1990’s, only a few individuals were partly to fully plated, and after 2000 all individuals were low plated. This illustrates a dramatic drop in plate number over time (Fig. 2). Because plate number evolution levels out when all individuals are low plated, this is most appropriately tested with a non-linear regression. We confirmed that plate number exhibits a significant exponential decay towards an asymptotic value (estimates: a = 1.02 [s.e.=0.669], b = 13.3 [s.e.=0.85; P < 0.0001], c = 0.085 [s.e. = 0.016; P = 0.007]). Over the study period, the stickleback became longer, increasing from a mean of 35.98 mm in 1974 to 46.54 mm in 2021 (Fig S1; t = 8.54, P < 0.0001), and increased in mass, from an average of 0.72 grams in 1974, to 1.89 grams in 2021 (Fig. S2; t = 7.79, P < 0.0001).

**Figure 2.**
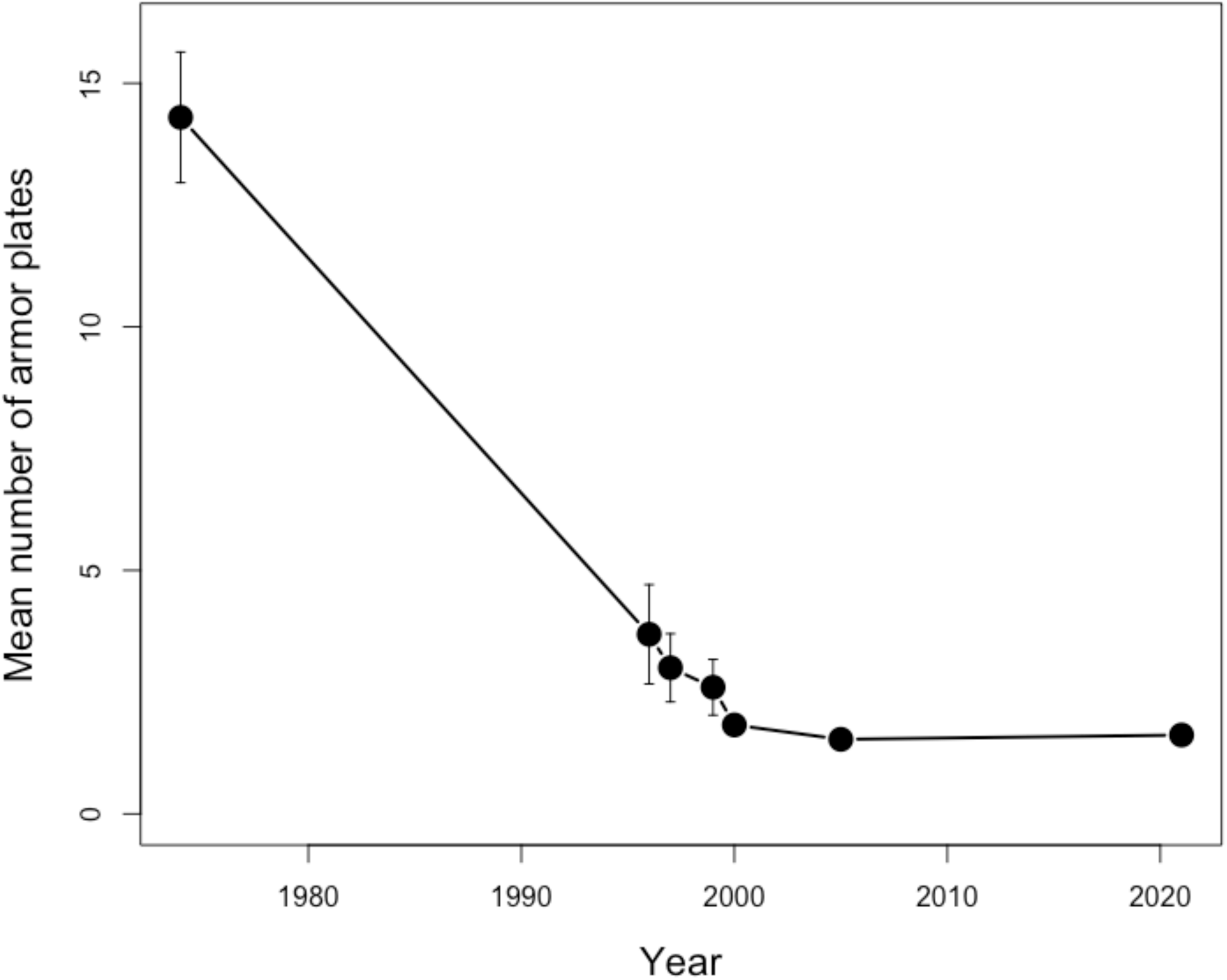
Mean armor plating over time, with circles representing mean values, vertical lines representing standard error. The armor plating evolution in this population follows the well-established trend of decreasing rapidly over a small number of generations to an average 1-2 plates on each side of the fish.

In addition to these well-known changes in morphology there were also major changes in the cestode infection rate. As expected, the newly established population was heavily infected. In 1974, eight years after the initial introduction, 71.4% of the stickleback were infected (95% CI 58%-83%). The prevalence of *S. solidus* tapeworms declined significantly over the 50 year period (Fig. 3A, binomial GLM z = -7.81, P < 0.0001). During this decline there was some year-to-year variation in the late 1990’s: infection rates rebounded from a low prevalence of 13% and 15% in 1997 and 1997, to 32% in 1999, before dropping to 5% in 2000. By 2021 we observed zero infections (0%, CI 0% - 12.7%). A similar decline was observed for infection intensity through time (Poisson GLM, z = -9.45, P < 0.0001).

**Figure 3.**
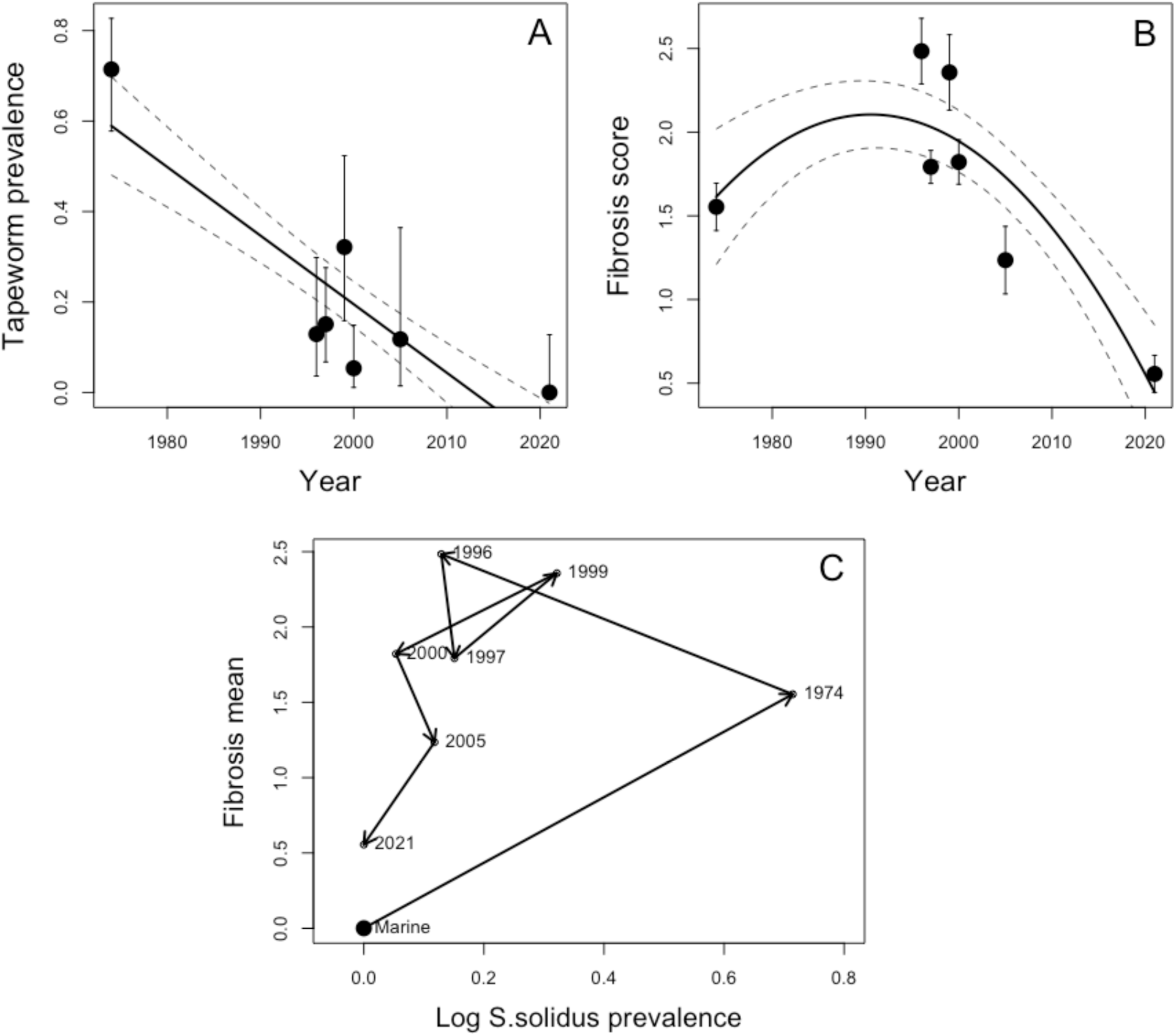
Tapeworm prevalence (A) and fibrosis score (B) over time. Tapeworm prevalence reduced dramatically over time, whereas fibrosis score increased and then decreased. The prevalence of *S. solidus* plotted with the mean fibrosis score (C) shows the combination of the two patterns: infection rates initially increase in a population with only modest fibrosis, then fibrosis increases and infection rates decrease, then both decrease dramatically. To represent the inferred ancestral state of marine fish, we add a large solid point (labeled ‘Marine’ with infection and fibrosis rate data from a nearby marine population (Sayward Estuary) collected in 2022.

Concurrent with the changes in tapeworm prevalence, we observed changing fibrosis through time (Fig. 3B). Compared to the initial mean fibrosis score of 1.55 in 1974, by 1996 fibrosis had increased by 60% to a high of 2.50, after which fibrosis again declined to a low of 0.56 in 2021 (a 78% decline from its high in 1996). Reflecting this transient increase in fibrosis, a quadratic model fit significantly better than a simpler linear model (linear t = 6.9, P < 0.0001; quadratic t = -7.0, P < 0.0001).

These changes in fibrosis were associated with changes in infection prevalence. In 1974, when the stickleback population would still be dominated by susceptible marine genotypes, infection rates were high and fibrosis severity moderate (Figure 3C). Two decades later, fibrosis was much stronger and infection prevalence lower, consistent with an evolutionary gain of fibrosis response conferring some protection against infection.

Parasite infection rate fell further in the 2000’s, and fibrosis rates fell accordingly. As tapeworm infections became rarer, fewer stickleback may have ingested infected copepods, and thus fewer individuals were stimulated to initiate fibrosis. By 2021, there were no detectable tapeworms present and few fish had been stimulated to initiate fibrosis. Overall across the whole time span, there was a non-significant correlation between annual mean fibrosis and parasite prevalence (r = 0.614, P = 0.1421). However, a linear test may not be suitable if the fibrosis-infection relationship is driven by two interrelated processes, the evolutionary gain of fibrosis response during high exposure risk, and a later phase of parasite decline, closely fitting our simple eco-evolutionary model (compare Fig. 1 to Fig. 3C). If we instead use a quadratic regression (favored by stepwise AIC, dAIC = 3.2), we find a nearly significant non-linear relationship between parasite prevalence and fibrosis (t = -2.4, P = 0.064). This nonlinear trend is consistent with an interpretation that fibrosis peaked at intermediate parasite prevalences during the collapse of the local *S. solidus* population.

Prior studies (of other populations showed that fibrosis severity differs heritably between populations (Weber et al. 2022, Hund et al. 2021, Bolnick et al. 2024). Therefore, changes in fibrosis levels may reflect evolutionary changes in the Heisholt population. If fibrosis response had indeed evolved during the span of our study, we would expect to find that initially, stickleback are unable to initiate a fibrosis response when infected, but in later years they do initiate fibrosis when infected. Fibrosis evolution should manifest as a significant year*infection interaction e_ect. Consistent with experimental evidence that infection induces fibrosis, tapeworm infection and fibrosis are positively related in the dataset overall (Binomial GLM, Z = 3.27, P = 0.0011), but this relationship changes with successive years (Z = -3.29, P = 0.0010). Specifically, at the outset of the Heisholt Quarry population, the mostly marine fish had low fibrosis regardless of infection status (Fig. 4, Fig. S3). In contrast, fibrosis was significantly related to infection status in the late 1990’s. This change in fibrosis-infection relationship is consistent with an evolutionary gain of fibrosis response to infection. However, the direction of the relationship is negative: fish with stronger fibrosis had fewer tapeworms. This negative relationship is consistent with a scenario in which infection induces fibrosis in some fish genotypes, which contributes to parasite clearance but fibrosis persists thereafter (as seen in laboratory infections). In later years (2005, 2021), tapeworms were scarce enough that we cannot effectively test for fibrosis-infection relationships.

**Figure 4.**
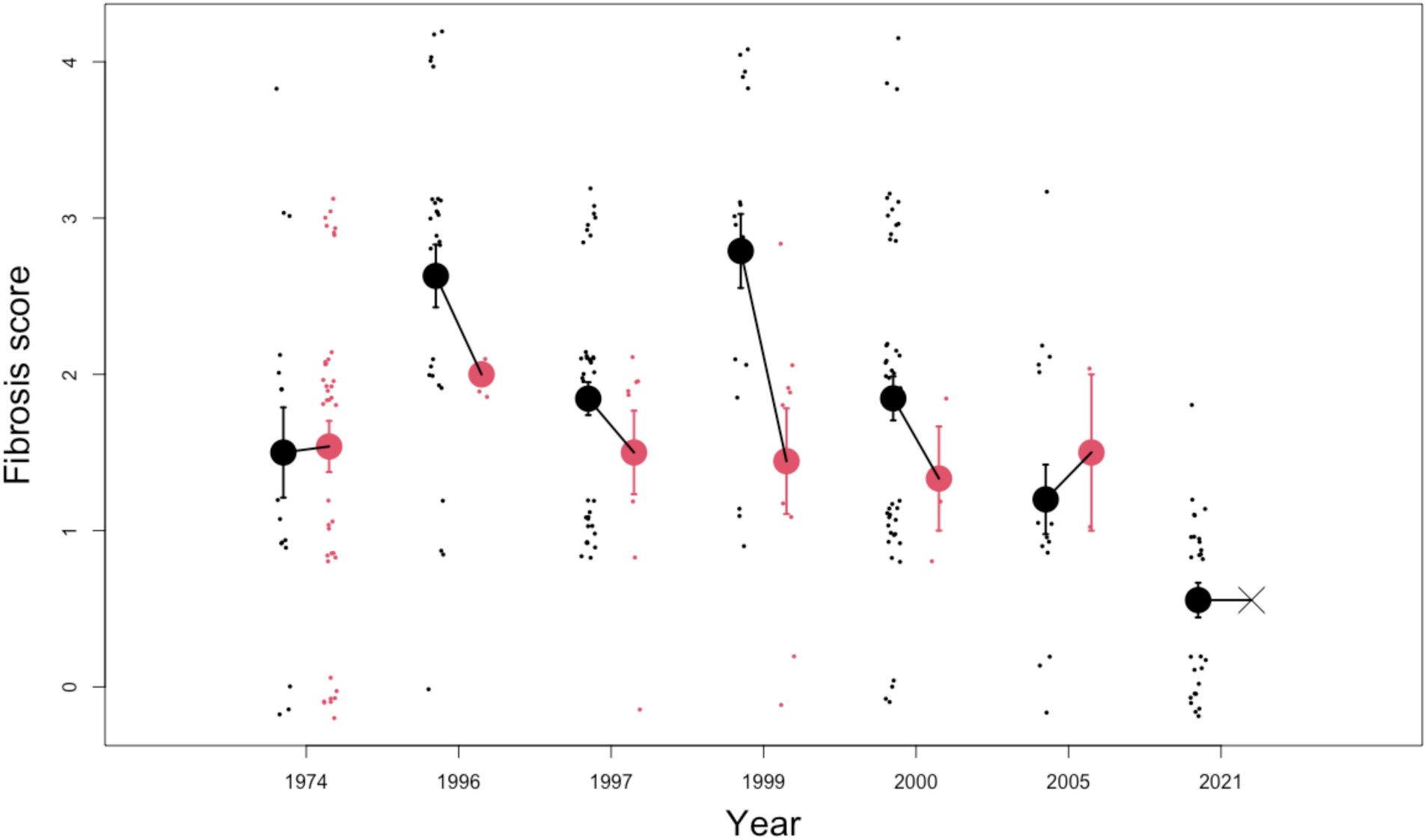
Fibrosis score of infected (red) and uninfected (black) fish. Large circles indicate mean value. Vertical lines indicate standard error. Diagonal lines indicate difference between infected and uninfected means. Small dots indicate individual data points (jittered for visualization). X indicates lack of data.

There was no significant change in log cestode mass over time (P > 0.1 for effects of year, infection intensity, and log fish mass). Stepwise AIC model comparisons favored retention of a model with infection intensity and fish mass, excluding year (deltAIC = 2.2), but even in the simplified model no term was significant.

## DISCUSSION

The experimentally founded stickleback population of Heisholt Quarry provides an opportunity to track eco-evolutionary changes in infection and immunity across a half century. Marine stickleback were introduced to the quarry in 1967, and sampled periodically between 1974 and 2021. Over this time span, the population evolved reduced armor plating, typical of newly established freshwater stickleback populations elsewhere (Bell et al. 2004; Colosimo et al. 2005). Selection pressures driving armor loss are very strong (Le Rouzic et al. 2011). However, we know far less about the parasitological changes and immunological adaptations during sticklebacks’ adaptation to freshwater. Indeed, there are few studies of vertebrates revealing the joint long-term dynamics of infection and immune traits in the wild (e.g., Bonneaud et al. 2018; Graham et al. 2010).

Previous work has established that marine stickleback are highly susceptible to *S.solidus* tapeworm infection (Weber et al. 2017a). They do not initiate a fibrosis response when exposed to live *S.solidus* (Weber et al. 2022), or injected with *S.solidus* antigens (Hund et al. 2021). When exposed to *S.solidus* in the lab, marine fish are infected at a high frequency and the parasites grow rapidly. This susceptibility reflects a lack of selection for resistance to *S.solidus* in marine environments, where *S.solidus* is absent. Upon colonizing Heisholt Quarry, susceptible marine fish would have been exposed to, and readily infected by, *S.solidus* which can disperse from nearby lakes via piscivorous birds. We indeed found that infection rates were high in the early 1974 sample from this lake. This finding matches observations in other recently-founded stickleback populations with marine immigrants, such as Cheney Lake in Alaska where infection prevalences can reach 100%.

Fibrosis is a plastic response to infection, being absent or minimal unless fish are stimulated by *S.solidus* antigens via live infection, or antigen injection (Hund et al. 2021). However, the capacity to initiate fibrosis is heritable. In an F2 hybrid cross between a high and low fibrosis population, a majority of fibrosis variance could be attributed to a large effect locus, *spi1b* (Weber et al. 2022). In common-garden reared fish from several populations in Alaska, tapeworm infection induced stronger fibrosis in fish descended from high-fibrosis wild populations (Bolnick et al. 2024b).

We expected that the initial colonists would be unable to initiate a fibrosis response, being dominated by non-fibrotic marine genotypes. Indeed, in 1974, stickleback exhibited low fibrosis whether or not they were infected (Fig. 3). But, in later generations we expected stickleback would evolve a capacity to initiate fibrosis when exposed to *S.solidus* (Fig. 1). By the late 1990’s, three decades after the marine fish were introduced, we found a substantial increase in fibrosis, and decline in parasite prevalence (Fig. 3). We also see a change in the fibrosis-infection relationship, indicating evolution of the response (Fig. 4, Fig. S5). Specifically, fish with stronger fibrosis were less likely to have a successful parasite infection. This is consistent with experimental evidence that fibrosis can lead to successful elimination of *S.solidus,* but fibrosis can linger for as long as a year after the immune challenge. Conversely in individuals that still lack the genetic capacity for fibrosis, we would observe successful tapeworms and low fibrosis scores.

The inferred evolution of fibrosis response in Heisholt Quarry coincided with the steady decline in *S.solidus* infection rates. The fibrosis immune response would reduce parasite reproductive rates if hosts successfully eliminated infections. But, fibrosis can also suppress parasite growth. *S.solidus* must reach a threshold mass in order to reproduce in birds, and above that threshold fecundity is positively related to mass (Lüscher and Wedekind 2002). Therefore, growth suppression in fibrotic fish could additionally contribute to reduced reproductive rate of the tapeworm population. By 2021, we detected no tapeworms at all in our sample, suggesting that stickleback immunity had been sufficient to eliminate the resident population (though immigration via bird dispersal would have continued). Without a viable resident population of tapeworms to induce an immune response, fibrosis rates were low. We speculate that this reduction in fibrosis is a loss of stimulation (low parasite exposure), rather than a genetic change in the host. However, there is evidence that fibrosis is costly (de Lisle et al. 2021, Weber et al. 2022, Matthews et al. 2024), and genetic loss of fibrosis has been documented in other populations (Weber et al. 2022). The potential for continued reintroduction of *S.solidus* from other populations would presumably maintain some resistance in the Heisholt Quarry population.

Cestode mass was not correlated with sample year, infection intensity, or fish mass. This is expected. As previous studies have noted, the peritoneal fibrosis response acts as an anti-cestode defense by restricting cestode growth (Weber et al. 2022), and sometimes killing cestodes. In the population studied here, most fibrotic fish were found without cestodes, suggesting that cestode death may be typical (which would obscure growth suppression effects).

Our results highlight the potential for newly established populations to encounter unfamiliar parasites and subsequently evolve increased immunity. In Heisholt Quarry, sticklebacks’ fibrosis response changed over a similar time scale as their loss of armor plating. This change is likely to be an evolutionary one, given previous evidence that the fibrosis response is heritable (Vrtilek and Bolnick 2021). Future work could confirm this inference through genomic analyses of the Heisholt Quarry population, or quantitative genetic study of F2 hybrids between Heisholt and marine fish. If this is indeed evolution, such a rapid shift in fibrosis would require quite strong selection, similar to the selection pressures existing for armor plate loss. Such selection for fibrosis is especially noteworthy given that fibrosis is widely viewed as an immune pathology in humans (Wynn 2008).

## ACKNOWLEDGEMENTS

We thank Mackenzie Kinney, & Stephanie Blaine for collecting samples in 2021, James MacLean and the late Don McPhail for samples from 1974, and Steven Vamosi for samples from the 90s and 2005. We thank Heather Alexander and Emma Choi for their comments and revisions to our manuscript. We thank Andrea Roth, Maria Rodgers, and Arshad Padhiar for their help with fibrosis scoring. This work was funded by NIAID grant 2R01AI123659-07 to DIB.

## AUTHOR CONTRIBUTIONS

DB and PS designed the study. DS collected some of the samples and provided all samples to DB and PS. PS dissected the fish and collected data. PS and DB analyzed the data, created figures, and wrote the original draft of the manuscript.

## DATA ARCHIVING

Data and R code to recreate statistical inferences and figures are archived on FigShare.org, at DOI: 10.6084/m9.figshare.28342601

## SUPPLEMENTARY MATERIALS

**Figure S1.**
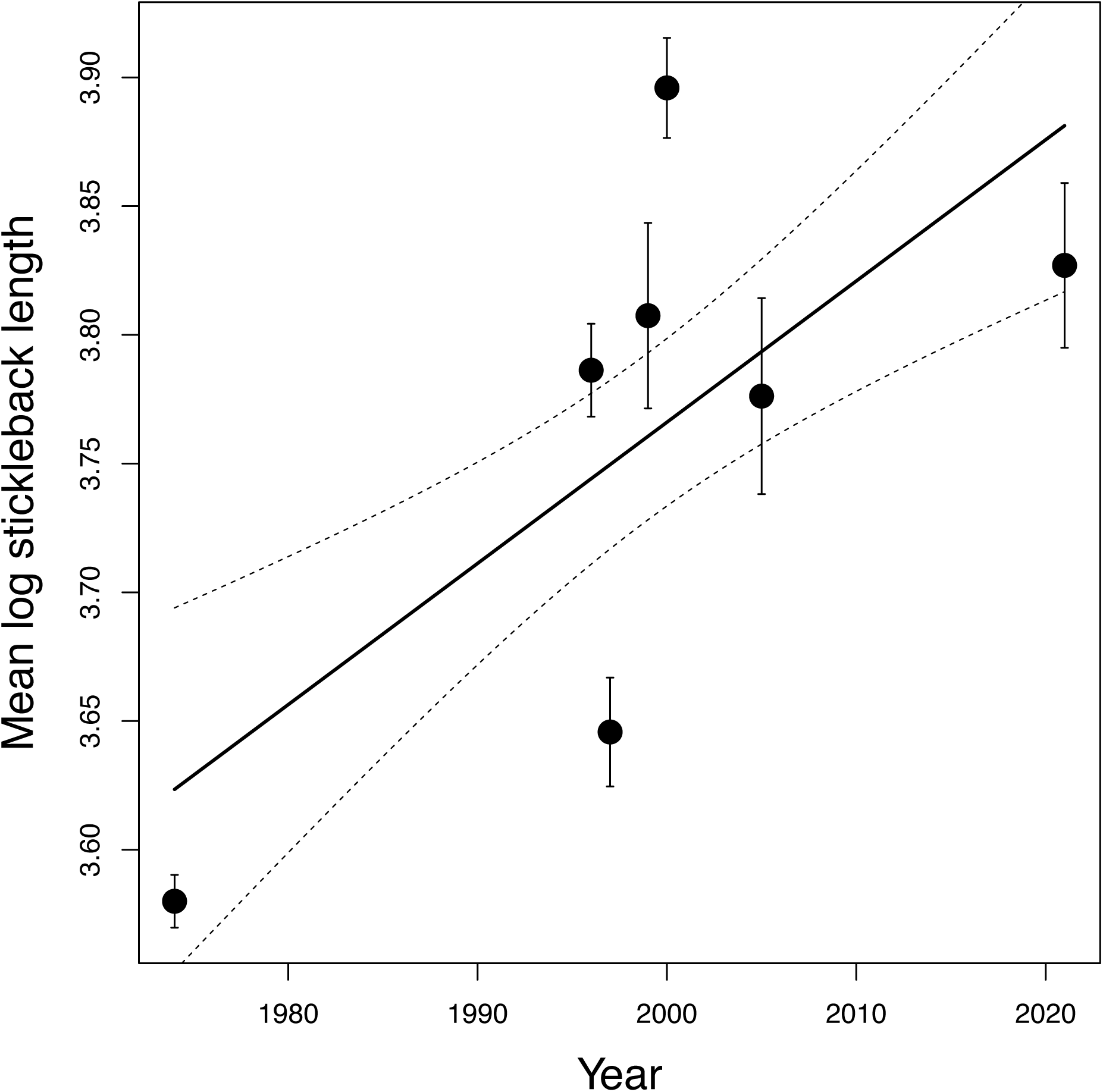
Log of average stickleback length each year. Stickleback on average got longer over time. Solid line indicates line of best fit, dotted line indicates standard error.

**Figure S2.**
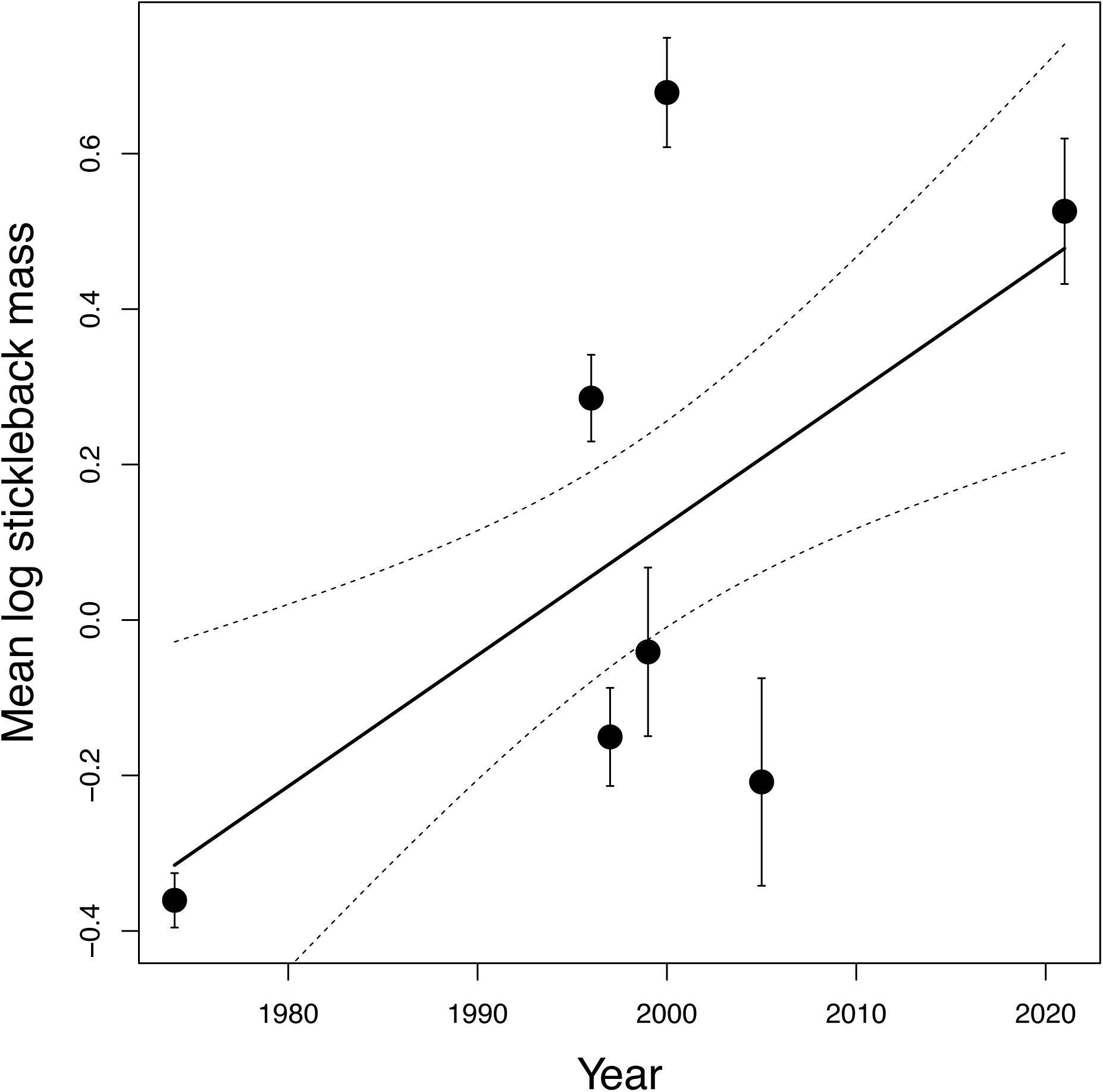
Log of average stickleback mass each year. Stickleback on average got larger over time, despite noticeable dips such as 2005. Solid line indicates line of best fit, dotted line indicates standard error.

**Figure S3.**
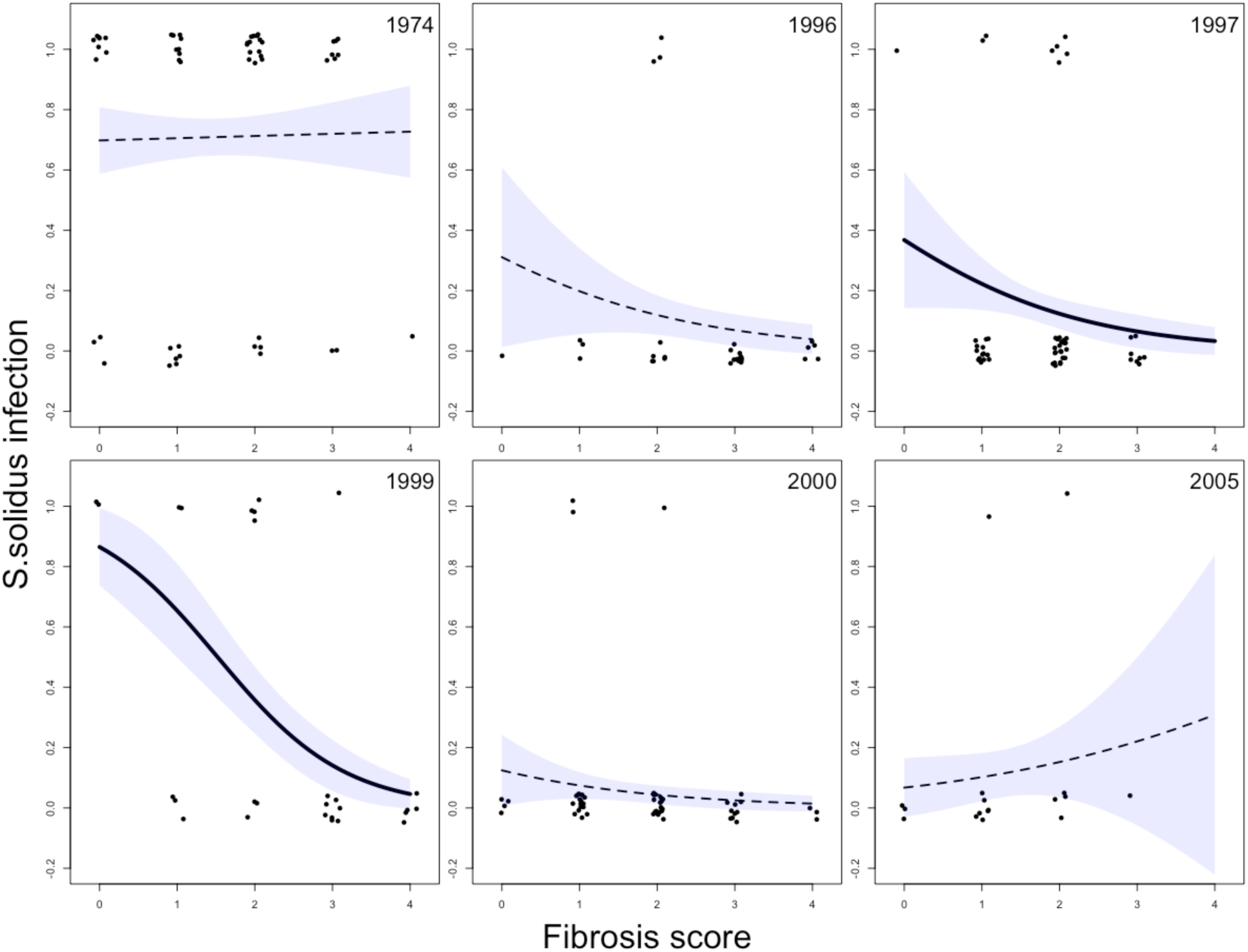
Fibrosis and cestode infection rates for six of the seven years included in the study (2021 excluded as there were no infections that year). Dots indicate individual fish with fibrosis score and infection (0 = not infected, 1 = infected), jittered for visualization. Line indicates general trend (logarithmic), shading indicates standard error.

## Notes

https://figshare.com/s/8e2cc46b82f49a215607

### Competing Interest Statement

The authors have declared no competing interest.

## LITERATURE CITED

Aguirre, W. E., K. Reid, J. Rivera, D. C. Heins, K. R. Veeramah, and M. A. Bell. 2022. Freshwater colonization, adaptation, and genomic divergence in threespine stickleback. Integrative and Comparative Biology 62(2):388–405.

Barber, I, and J. P. Scharsack. 2009. The three-spined stickleback-*Schistocephalus solidus* system: an experimental model for investigating host-parasite interactions in fish. Parasitology 137:411–424.

Bell, M. A., W. E. Aguirre, and N. J. Buck. 2004. Twelve years of contemporary armor evolution in a threespine stickleback population. Evolution 58(4):814–824.

Bell, M. A., D. C. Heins, M. A. Wund. F. A. von Hippel, R. Masenglil, K. Dunker, G. A. Bristow, and W. E. Aguirre. 2016. Reintroduction of Threespine stickleback into Cheney and Scout lakes, Alaska. Evolutionary Ecology Research 17:157–178.

Bolnick, D.I., S. Arruda, C. Polonia, L. Simonse A. Padhiar, A. Roth, M. Rodgers. 2024a. The dominance of coinfecting parasites’ indirect genetic effects on host traits. American Naturalist. 204: 482–500.

Bolnick, D.I. A.P. Hendry, C.L. Peichel, R.D.H. Barrett, C. Wolf, T. Sasser, E. Kearns, N. Steinel, J. Weber. 2024b. Destabilized host-parasite dynamics following the founding of new populations. BioRxiv preprint: https://biorxiv.org/cgi/content/short/2024.06.24.600494v1

Bonneaud, C., S. L. Balenger, A. F. Russel, J. Zhang, G. E. Hill, and S. V. Edwards. 2011. Rapid evolution of disease resistance is accompanied by functional changes in gene expression in a wild bird. PNAS 108(19):7866–7871.

Bonneaud, C., M. Giraudeau, L. Tardy, M. Staley, G.E. Hill, and K.J. McGraw. 2018. Rapid antagonistic coevolution in an emerging pathogen and its vertebrate host. Current Biology. 28:2978–2983.

Colosimo, P. F., K. E. Hosemann, S. Balabhadra, G. Villareal Jr., M. Dickson, J. Grimwood, J. Schmutz, R. M. Meyers, D. Schluter, and D. M. Kingsley. 2005. Widespread parallel evolution in sticklebacks by repeated fixation of ectodysplasin alleles. Science 307:1928–1933.

Confer, A., V. Vu, C. J. Drevecky, and W. E. Aguirre. 2012. Occurrence of *Schistocephalus solidus* in anadromous threespine stickleback. Journal of Parasitology 98(3): 676–678.

Crossan, J., S. Paterson, and A. Fenton. 2007. Host availability and the evolution of parasite life-history strategies. Evolution 61(3):675–684.

De Lisle, S. P., and D. I. Bolnick. 2021. Male and female reproductive fitness costs of an immune response in natural populations. Evolution 75(10):2509–2523.

Ebert, D. 2008. Host-parasite coevolution: insights from the Daphnia-parasite model system. Current Opinion in Microbiology 11:290–301.

Graham, A.L., A.D. Hayward, K.A. Watt, J.G. Pilkington, J.M. Pemberton, and D.H. Nussey. 2010. Fitness correlates of heritable variation in antibody responsiveness in a wild mammal. Science. 330:662–665.

Hund, A. K., L. E. Fuess, M. L. Kenney, M. F. Maciejewski, J. M. Marini, K. C. Shim, and D. I. Bolnick. 2022. Population-level variation in parasite resistance due to differences in immune initiation and rate of response. Evolution Letters 6(2):162–177.

Kirch, M., Romundset, A., Gilbert, M.T.P., Jones, F.C., and A.D. Foote. 2021. Ancient and modern stickleback genomes reveal the demongrahic constraints on adaptation. Current Biology. 31:2027–2036.

Kristjánsson, B. K., S. Skúlason, and D. L. G. Noakes. 2002. Rapid divergence in a recently isolated population of threespine stickleback (*Gasterosteus aculeatus* L.). Evolutionary Ecology Research 4:659–572.

Lachish, S., S. C. L. Knowles, R. Alves, M. J. Wood, and B. C. Sheldon. 2011. Fitness effects of endemic malaria infections in a wild bird population: the importance of ecological structure. Journal of Animal Ecology 80:1196–1206.

Le Rouzic, A., K. Ostbye, T. O. Klepaker, T. F. Hansen, L. Bernatchez, D. Schluter, and A. Vollestad. 2011. Strong and consistent natural selection associated with armour reaction in sticklebacks. Molecular Ecology 20:2483–2493.

Lohman, B. K., N. C. Steinel, J. N. Weber, and D. I. Bolnick. 2017. Gene expression contributes to the recent evolution of host resistance in a model host/parasite system. Frontiers in Immunology 8:1071.

Lüscher, A., and C. Wedekind. 2002. Size-dependent discrimination of mating partners in the simultaneous hermaphroditic cestode *Schistocephalus solidus*. Behavioral Ecology 13(2):254–259.

Jones, F. C, M. G. Grabherr, Y. F. Chan, P. Russell, E. Mauceli, J. Johnson, R. Swofford, M. Pirun, M. C. Zody, S. White, E. Birney, S. Searle, J. Schmutz, J. Grimwood, M. C. Dickson, C. T. Miller, B. R. Summers, A. K. Knecht, S. D. Brady, H. Zhang, A. A. Pollen, T. Howes, C. Amemiya, Broad Institute Genome Sequencing Platform and Whole Genome Assembly Team, E. S. Lander, F. Di Palma, K. Linblad-Toh, and D. M. Kingsley. 2012. The genomic basis of adaptive evolution in threespine sticklebacks. Nature 484:55–61.

Maclean, J. A. 1974. Variation and Natural Selection in a Population of Sticklebacks (*Gasterosteus)*. [Doctoral thesis, University of British Columbia] 10.14288/1.0099961

MacLean, J. 1980. Ecological genetics of threespine sticklebacks in Heisholt Lake. Canadian Journal of Zoology 58:2026–2039.

MacDonald, H., and D. Brisson. 2022. Host phenology regulates parasite-host demographic cycles and eco-evolutionary feedbacks. Ecology and Evolution 12(3):e8658.

Matthews, D. G., M. F. Maciejewski, G. A. Wong, G. V. Lauder, and D. I. Bolnick. 2023. Locomotor e_ects of a fibrosis-based immune response in stickleback fish. Journal of Experimental Biology 226(23): jeb246684.

Nordeide, J. T., and F. Matos. 2016. Solo Schistocephalus solidus tapeworms are nasty. Parasitology 143(10):1301–1309.

Nørgaard, L. S., B. L. Phillips, and M. D. Hall. 2019. Infections in patchy populations: Contrasting pathogen invasion success and dispersal at various times since host colonization. Evolution Letters 3(5):555–566.

Orti, G., M. A. Bell, T. E. Reimchen, and A. Meyer. 1994. Global survey of mitochondrial dna sequences in the threespine stickleback: evidence for recent migrations. Evolution 48(3):608–622.

Pelletier, F., Garant, D., & Hendry, A. P. (2009). Eco-evolutionary dynamics. Philosophical Transactions of the Royal Society B: Biological Sciences, 364:1483–1489.

R Core Team. 2023.R: a language and environment for statistical computing. R. foundation for statistical computing, Vienna, Austria.

Simmonds, N. E., and I. Barber. 2016. The effect of salinity on egg development and viability of *Schistocephalus solidus* (cestoda: diphyllobothriidea). Journal of Parasitology 102(1):42–46.

Thompson, R. C. A. and A. J. Lymbery. 1996. Genetic variability in parasites and host-parasite interactions. Parasitology 112:S7:S22.

Turola, E., S. Petta, E. Vanni, F. Milosa, L. Valenti, R. Critelli, L. Miele, L. Maccio, V. Calvaruso, A. L. Fracanzani, M. Bianchini, N. Raos, E. Bugianesi, S. Mercorella, M. Di Giovanni, A. Craxì, S. Fargion, A. Grieco, C. Camma, F. Cotelli, and E. Villa. 2015. Ovarian senescence increases liver fibrosis in humans and zebrafish with steatosis. Disease Models and Mechanisms 8:1037–1046.

Vamosi, S. M. 2006. Contemporary evolution of armour and body size in a recently introduced population of threespine stickleback *Gasterosteus aculeatus*.

Vrtilek, M., and D. I. Bolnick. 2021. Macroevolutionary foundations of a recently evolved innate immune defense. Evolution 75(10):2600–2612.

Weber, J. N., M. Kalbe, K. C. Shim, N. I. Erin, N. C. Steinel, L. Ma, and D. I. Bolnick. 2017a. Resist Globally, infect locally: a transcontinental test of adaptation by stickleback and their tapeworm parasite. The American Naturalist 189(1):43–57.

Weber, J. N., N. C. Steinel, K. C. Shim, and D. I. Bolnick. 2017b. Recent evolution of extreme cestode growth suppression by a vertebrate host. PNAS 114(25) 6575–6580.

Weber, J. N., N. C. Steinel, F. Peng, K. C. Shim, B. K. Lohman, L. E. Fuess, S. Subramanian, S. P. De Lisle, and D. I. Bolnick. 2022. Evolutionary gain and loss of a pathological immune response to parasitism. Science 377(6611):1206–1211.

Wynn, T. 2008. Cellular and molecular mechanisms of fibrosis. Journal of Pathology. 214:199–210.

